# Grip Force Complexity Under Constraint: Evidence from Fractal and Entropy Analyses

**DOI:** 10.64898/2026.05.31.729065

**Authors:** Louis Cointre, Fabien Buisseret, Frédéric Dierick, Nicolas Boulanger, Olivier White

## Abstract

The present study investigates how grip force (GF) and load force (LF) dynamics reorganize under varying task constraints, focusing on the fractal and entropic properties of motor output. Twenty healthy adults performed precision grip tasks across five force conditions: two spontaneous conditions (pre, post) without visual feedback and three target-driven conditions (natural, −10%, +10% of natural). The temporal and informational structure of GF and LF signals were quantified using the Hurst exponent (H) and Sample Entropy (SampEn), capturing long-range temporal organization and local irregularity. Constrained conditions reduced GF coefficient of variation but increased both H and SampEn relative to spontaneous pre-trials, while LF indices remained largely unchanged. Thus, grip control under constraint became more temporally persistent and locally irregular, suggesting a more structured temporal organization rather than a simple loss of complexity. Intra-trial analyses further revealed an increase in H from the first to the second half of spontaneous trials, consistent with progressive self-organization in the absence of explicit constraints. Across conditions, H and SampEn were positively correlated for both GF and LF, suggesting that predictability and complexity are not necessarily inversely related in this context. Overall, these findings suggest that the human motor system adapts to force constraints not by suppressing variability, but by reorganizing it across scales, combining temporal persistence with local flexibility. This multidimensional characterization of variability may help refine theoretical models of optimal movement variability and may inform clinical or training approaches aimed at assessing or enhancing neuromotor adaptability.

## Introduction

Grasping is a fundamental component of human motor control, playing a central role in the execution of everyday actions. Since the pioneering work of Napier (1956), it has been widely acknowledged that grasping primarily relies on two types of grips: the precision grip and the power grip. Each involves distinct neuromuscular strategies to modulate the applied force according to the properties of the object and the environmental context (Napier, 1956). This adaptive capacity relies on coordinated sensorimotor mechanisms involving cutaneous and proprioceptive receptors, such as Meissner’s corpuscles, Merkel cells, Pacinian corpuscles, and Ruffini endings, as well as muscle spindles and Golgi tendon organs. These receptors continuously provide the central nervous system with information on texture, pressure, vibration, and muscle tension, thereby enabling the fine regulation of grasping (Johansson & Flanagan, 2009; Proske & Gandevia, 2012).

Grip force (GF) is essential for stabilizing an object, preventing it from slipping or being crushed. Its dynamic modulation depends on the integration of multiple sensory inputs and the regulation of a “safety margin”, defined as the excess force applied beyond the minimum required to ensure object stability (Danion, 2008; Westling & Johansson, 1984). In normal conditions, the safety margin amounts to about 20% but can increase dramatically—sometimes to several hundred percent—in certain pathologies (Hermsdörfer et al., 2003; Lodha et al., 2012). Human force output is never perfectly constant, even when the task requires maintaining a steady level. This variability, once dismissed as mere motor noise, is currently recognized as a meaningful marker of the adaptive complexity of motor control (Stergiou et al., 2006; Vaillancourt & Newell, 2002). Indeed, traditionally, motor variability was regarded as unwanted noise—an error to be minimized in the pursuit of stable and accurate performance. Growing evidence across physiology, neuroscience, and cognitive science demonstrates that variability is not merely random noise but a functional property of living systems. It emerges from nonlinear interactions within complex networks and reflects the system’s ability to adapt, self-organize, and remain flexible under uncertainty (Goldberger, 1991; Goldberger, Amaral, et al., 2002). Rather than degrading performance, variability can enhance adaptability, resilience, and health: low levels of noise can promote flexibility (Casartelli et al., 2023), stochastic resonance has been shown to improve posture and balance (White et al., 2018), and fractal or chaotic fluctuations have been identified as prognostic markers in diverse conditions ranging from gait dysfunction (Dierick et al., 2021; Hausdorff, 2007) to cardiovascular health (Gronwald et al., 2020), epilepsy (Li et al., 2005), mood disorders (Akar et al., 2015), and even artistic performance (Robles et al., 2021). Faisal and colleagues highlighted that neuromotor noise actively contributes to this adaptive flexibility by enabling the exploration of multiple motor solutions (Faisal et al., 2008). Despite this broad evidence, such a perspective has rarely been applied to grip force, a fundamental component of human action.

Among nonlinear analyses, fractal approaches have been proposed to characterize this complex organization. Biological processes, including GF, exhibit self-similar properties across multiple temporal scales—reflecting a highly organized adaptive system. This has been repeatedly demonstrated by Goldberger and colleagues (Goldberger, 1991, 1996, 1997; Goldberger et al., 1990; Goldberger, Peng, et al., 2002). The Hurst exponent (H), estimated through Detrended Fluctuation Analysis (DFA), provides a robust index of the temporal persistence of force fluctuations (Peng et al., 1995). When H approaches 0.5, the signal tends toward random behavior; in contrast, higher values indicate long-range temporal persistence, reflecting a tendency for the signal to maintain its direction over time. Complementary to this fractal characterization, entropic measures capture a distinct dimension of signal complexity. Sample Entropy (SampEn) quantifies the local unpredictability of a time series by assessing the likelihood that patterns recurring over short epochs will remain similar at the next comparison—thus reflecting the short-term irregularity of the signal rather than its long-range structure (Richman & Moorman, 2000). Thus, H and SampEn capture complementary, rather than redundant, properties of the signal: H characterizes how force fluctuations are organized across time scales, while SampEn captures their local irregularity. In this study, we use the term *complexity* to designate the joint configuration of these two properties—long-range temporal persistence and local unpredictability—rather than treating complexity as a single scalar quantity.

The introduction of constraints, such as imposing target force ranges, can disrupt the spontaneous dynamics of motor control. The motor system’s ability to maintain an optimal level of complexity appears to be compromised under constraint. According to the “Optimal Movement Variability” model proposed by Stergiou and Decker, an excessive reduction in adaptive variability may reflect a loss of behavioral flexibility, making the system more vulnerable to perturbations (Stergiou & Decker, 2011). Fernandes and Chau showed that imposed force adjustment alters adaptive strategies, potentially affecting signal complexity. In this sense, constraints may promote a more rigid form of control, characterized by a reduction in both temporal and spatial complexity (Fernandes & Chau, 2008; Warlop et al., 2013). Despite growing interest in these issues, no studies have examined the effect of varying levels of constraint on GF complexity. It remains unknown whether varying the intensity of force constraints (below vs. above natural level) produces graded or qualitative changes in these dynamics.

Here, we examine how varying levels of force constraints influence the fractal dynamics of grip motor control. Specifically, we distinguish five conditions: two spontaneous conditions (pre and post), during which participants grip naturally without any explicit target; a natural condition, in which they are asked to reproduce their own spontaneous GF (i.e., the pre value); and two constrained conditions, requiring GF that are either 10% above (+10%) or below (–10%) the natural level with the help of a biofeedback system. This design enables us to evaluate how increasing constraint intensity progressively disrupts the system’s natural complexity. We hypothesized that, in the absence of explicit constraints, spontaneous GF production would exhibit a balanced temporal organization, characterized by a combination of long-range temporal persistence and local irregularity. In contrast, we expected externally imposed force targets to constrain the system and reduce this temporal richness, leading to lower SampEn and a shift of H toward more rigid dynamics. To test this prediction, we analyzed GF fluctuations using complementary indices (H and SampEn) across five force conditions to capture changes in both long-range temporal structure and local signal irregularity. The condition-based comparisons of H, SampEn, and CV were hypothesis-driven, with predictions derived from the Optimal Movement Variability framework; the inter-index correlation analyses were exploratory in nature and intended to characterize the multidimensional structure of motor variability across force conditions.

## Materials and Methods

### Participants

Twenty healthy right-handed participants (13 men, 7 women), aged between 20 and 28 years (mean ± standard deviation: 21 ± 2 years), took part in the study. Exclusion criteria included a recent (<6 months) history of traumatic injury or surgery of the upper limb, especially to the dominant hand and wrist (fractures, sprains) and any central or peripheral neurological disorders (e.g., multiple sclerosis, neuropathies) affecting fine motor control. The protocol complied with the Declaration of Helsinki and was approved by the French national ethics committee for research in physical activity (CERSTAPS, registration number: IRB00012476-2021-06-12-139, date of approval: 6 December 2021). Written informed consent was obtained prior to participation.

### Experimental Procedure and Apparatus

Participants were seated in a standardized position in front of a monitor, with their dominant arm positioned alongside the body, the elbow flexed at 90°, and the wrist relaxed. The experimental task required participants to grasp a custom-built object instrumented for force measurement. Grip forces were measured using a 6-axis Mini40 Force/Torque transducer (diameter: 40 mm; height: 12.2 mm; base mass of 50 g; ATI Industrial Automation, Apex, NC, USA). To standardize the experimental conditions and adjust the mechanical resistance, a 200-g counterweight was securely attached to the transducer assembly using the integrated mounting interface, resulting in a total instrumented object mass of 250 g. The transducer was interfaced with a BIOPAC MP160 data acquisition system. Force signals (F_x_, F_y_, F_z_) were digitized and recorded using AcqKnowledge software (v5.0.8.1) at a constant sampling frequency of 500 Hz.

Following the setup described above, the experimental task followed a structured sequence of grasp, lift, static hold, and release. Each trial, from the initial grasp to the final release, lasted for 45 seconds. The sequence was guided by specific verbal cues. Grasp and lift: Upon the command “Ready? You may grasp” (“*Prêt? Tu peux saisir*”), participants grasped the device using a precision grip and lifted it naturally approximately 20 cm off the table, ensuring the distal phalanges of the thumb and index finger remained parallel and the object was held vertically. Static Hold: Participants then adjusted their GF to match a designated target level. Release: The trial concluded with the command “You may set it down” (“*Tu peux poser*”).

The protocol was divided into three phases: two spontaneous conditions (pre and post) without visual feedback, and one constrained condition providing real-time visual feedback. In the constrained condition, the monitor displayed three target force levels represented by horizontal cursors (Fig. 1): natural force (reference) indicated by a black cursor, Low force (−10%) indicated by a blue cursor, and High force (+10%) indicated by a red cursor. The reference natural force was defined as the mean GF computed over the selected hold segments of all trials in the initial spontaneous pre-condition. It should be noted that in the constrained condition, the visual display served not as supplementary feedback on an otherwise unchanged task, but as the operational definition of the target itself: participants were required to actively regulate GF in real time to match an explicit force reference, constituting a qualitatively distinct motor behavior from unconstrained force maintenance. The −10% and +10% targets were then defined as ±10% of this participant-specific value. Participants performed 10 trials per condition in a randomized order, with 10 seconds of rest between trials. Any non-compliant trial, such as those involving incorrect force levels or postural deviations (e.g., involuntary arm movements or lifting the forearm), was immediately repeated. To prevent motor fatigue, additional rest periods were provided upon request. The entire session lasted approximately one hour.

**Figure 1.**
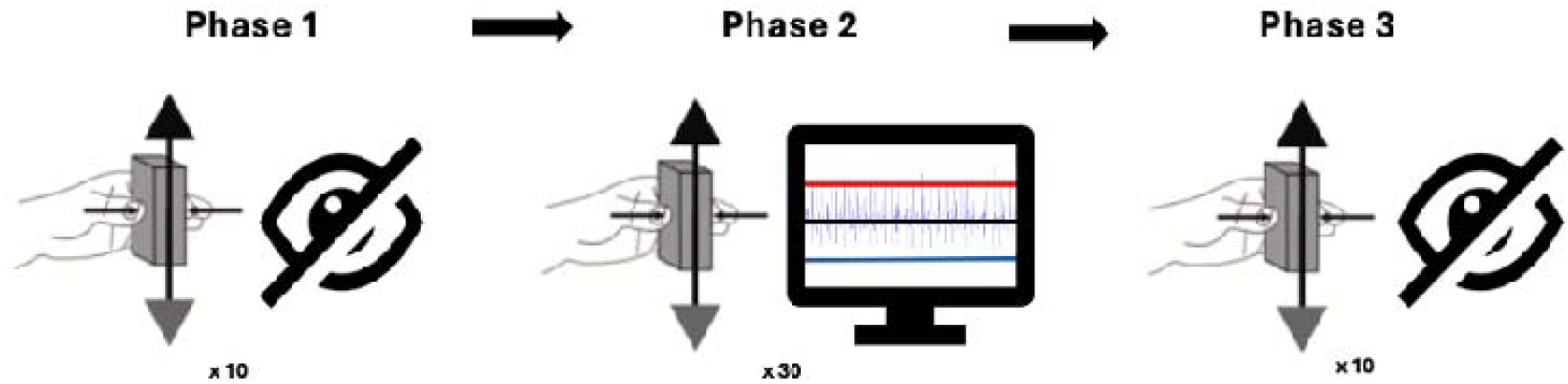
Illustration of the experimental procedure and its different phases. **Phase 1** (spontaneous pre): Participants performed 10 trials of static lifting and holding without visual feedback to establish their natural grip force baseline. **Phase 2**: (constrained): Participants performed 30 trials with real-time visual feedback on the monitor. They were instructed to match their GF to one of three randomized target cursors: +10% (red), natural reference (black), and −10% (blue). **Phase 3**: (spontaneous post): A final condition of 10 trials was performed without visual feedback to assess potential after-effects or fatigue. The icon with the cross-out eye denotes the absence of visual feedback. Vertical arrows indicate the lifting and holding instructions. Horizontal arrows illustrate GF.

### Data Processing

Grip force (GF) was defined as the force component normal to the sensor surface, with sign reversed (−Fz), whereas load force (LF) was computed as the Euclidean norm of the tangential components:—. Data were stored in Excel files. A digital low-pass zero phase lag Butterworth filter with a relatively high cutoff frequency (30 Hz, 8^th^ order) compared to what is usually adopted in similar experiments (White et al., 2005) was applied to remove high-frequency, non-physiological noise. Stable segments during the hold phase of the object were selected manually from each trial based on visual inspection. The initial grasping phase and the final release phase were systematically excluded, and a continuous segment of approximately 35 s (corresponding to 17 500 samples) was retained for analysis in each trial. Segment length was kept identical across trials and conditions.

We ran analyses to verify that the three imposed force level constraints differed significantly from one another. A Friedman test on participant-level mean GF revealed a strong overall effect of condition (*χ*^2^(2) = 937.57, p < 0.001, Kendall’s W = 0.56), confirming that participants modulated GF according to each target level (−10% vs natural: p = 0.022; −10% vs + 10%: p < 0.001; natural vs +10%: p = 0.050). Further comparisons between spontaneous conditions (pre and post) and constrained conditions revealed significant differences, particularly for the −10% and +10% levels. These results confirm the validity of the ±10% target ranges, symmetrically imposed around the natural force level to ensure consistency. This symmetry was designed to avoid biases in the imposed constraints, while accounting for a physiologically acceptable safety margin around spontaneous force production. It also allows the natural force to be bound without inducing excessive overload, while ensuring sufficient variation to detect significant condition-related effects.

The extracted indices characterized complementary aspects of force-control variability and were calculated separately on LF and GF. The coefficient of variation (CV), defined as the ratio of the standard deviation to the mean force, was computed to quantify relative variability. H was estimated using detrended fluctuation analysis (DFA) to assess the temporal predictability of the force signal, with values approaching 1 indicating stronger long-term dependencies. First-order DFA was applied to the mean-centered signal, using window sizes ranging from 3 to 316 samples (logarithmically spaced), and H was derived from the slope of the linear portion of the log F(n) vs. log n plot. SampEn was calculated for each time segment to evaluate local signal irregularity, using an embedding dimension of m = 2, a tolerance r = 0.2 × SD, and a time delay τ = 1. In addition, to capture potential intra-trial changes in temporal predictability, each trial was divided into two equal-length halves, and H was estimated separately for the first (H_early) and second (H_late) halves. For detailed mathematical definitions and theoretical justifications of these indices, the reader is referred to Peng et al. for H estimated via DFA and to Richman and Moorman for SampEn (Peng et al., 1995; Richman & Moorman, 2000).

### Statistical Analysis

Data preprocessing (filtering, averaging, feature extraction, visualization) was performed in Python using NumPy, Pandas, SciPy, and Seaborn libraries, along with the specific *hurst* and *hfda* modules. The dataset had a within-subject structure, with five repeated conditions per participant (spontaneous pre, −10%, natural, +10%, spontaneous post), and ten trials per condition. For each participant and condition, the ten trials were averaged to obtain a single value per variable, yielding 20 participant-level observations per condition.

Normality was tested using the Shapiro–Wilk test. As none of the variables met the normality assumption, all comparisons were conducted using non-parametric tests. Comparisons between mean GF and LF were performed using paired-sample Wilcoxon tests. The effect of force level on each variable was analyzed separately for GF and LF using Friedman tests applied to the participant-level means.

When significant effects were found, Wilcoxon signed-rank tests with Bonferroni correction were applied for post-hoc comparisons. The significance threshold was set at p < 0.05 for all analyses. Additional exploratory analyses examined the evolution of each variable across trials. No systematic trial-to-trial trends were observed for variability or entropy indices, and effects on force signal persistence were small across conditions.

Relationships between the extracted indices were explored using Spearman’s rank-order correlations (ρ), displayed as heatmaps. Correlations were computed on participant-level mean values across conditions, yielding 100 observations per force component (20 participants × 5 conditions) for each index. This method was chosen due to the non-normality of the distributions and the robustness of Spearman’s test for detecting monotonic relationships. All statistical analyses were conducted using JASP (v0.18.3). To limit the risk of Type I errors across the full analysis pipeline, confirmatory tests (Friedman and post-hoc Wilcoxon) were conducted separately for GF and LF with Bonferroni correction applied within each family of comparisons. The intra-trial comparison (H_early vs. H_late) constitutes a single planned contrast per condition and was not subject to further correction. Spearman correlation analyses were treated as exploratory and interpreted accordingly, without adjustment for multiple comparisons.

## Results

### Analysis of Complexity and Variability Indices

The use of the CV, H, and SampEn enables an exploration of motor variability across distinct yet complementary dimensions. CV quantifies the overall amplitude of fluctuations, independently of their temporal organization. In contrast, H evaluates the structure of temporal dependencies, revealing whether successive signal variations are correlated or independent—an important indicator of the persistence of motor control. SampEn, finally, captures the local complexity of the signal by assessing its irregularity or unpredictability over short time scales.

Analysis of the CV (Fig. 2A) revealed a significant main effect of force condition, *(4)* = 153.92, p < 0.001, Kendall’s W = 0.18, indicating a moderate effect size. Variability was consistently greater for *GF* than for LF. Post-hoc Wilcoxon signed-rank tests with Bonferroni correction showed that *GF* variability was significantly reduced under constrained conditions compared with spontaneous pre and post conditions (all p < 0.001), while no significant differences were found between constrained levels (p = 1.000). LF variability remained unchanged across conditions (all p = 1.000). A paired Wilcoxon test confirmed overall greater variability in *GF* (M = 0.0419, SD = 0.0324) than LF (M = 0.0251, SD = 0.0257; z = 12.383, p < 0.001).

**Figure 2.**
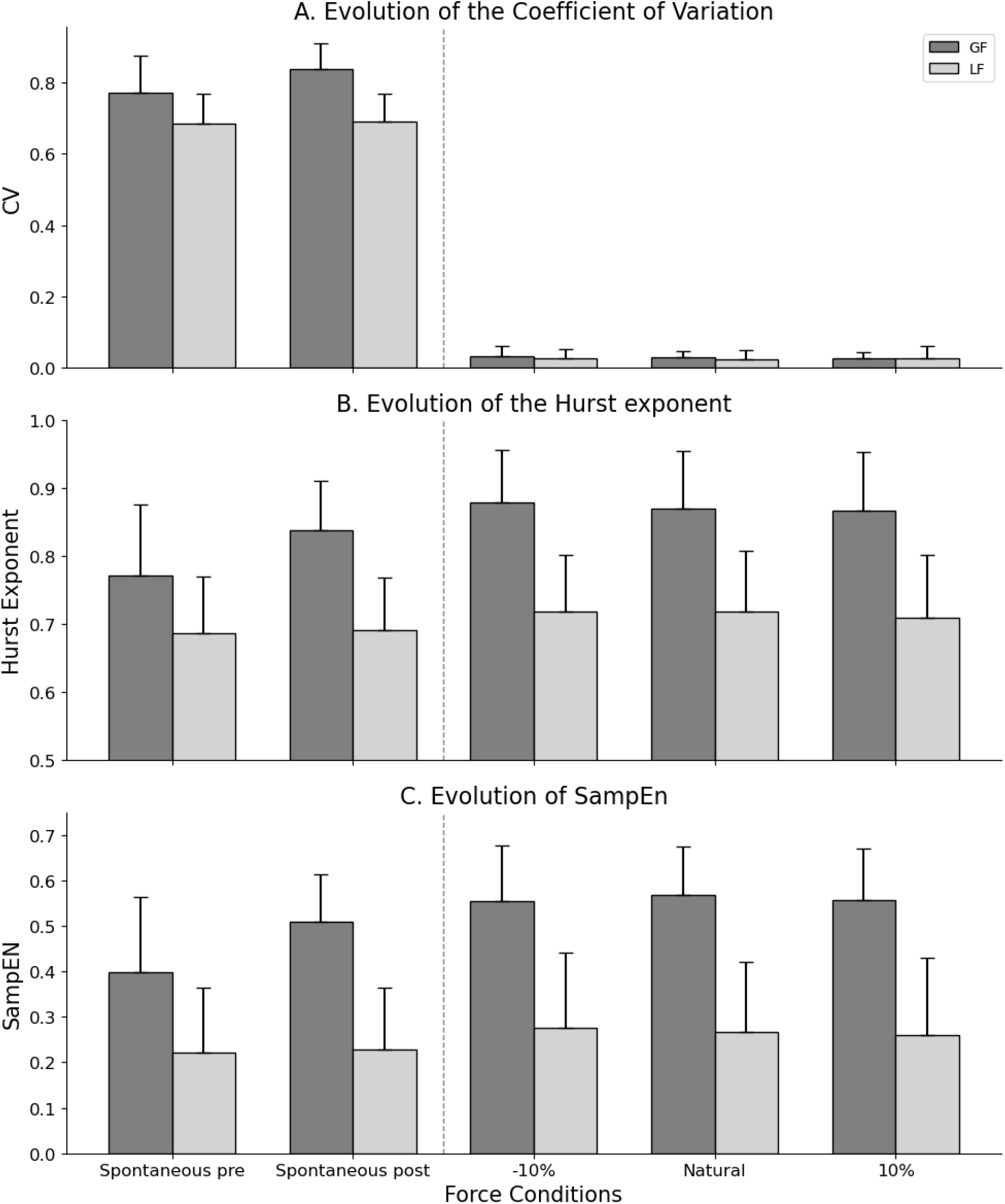
Variability indices of GF (dark grey) and LF (light grey) broken down into coefficient of variation (A), H (B) and SampEn (C) across the five force conditions.

H analysis (Fig. 2, B) revealed a robust effect of force condition, *χ*^2^ *(4)* = 668.05, p < 0.001, Kendall’s W = 0.79, indicating a very large effect size. Post-hoc Wilcoxon signed-rank tests with Bonferroni correction showed that, for *GF*, temporal predictability (H) was significantly higher in all constrained conditions compared with spontaneous pre and post conditions (all p < 0.001), and that H also differed significantly between the two spontaneous conditions (pre vs post: p < 0.001). No significant differences were observed among the constrained levels (all p = 1.000). No significant differences were observed for LF (all p > 0.05). Finally, a paired Wilcoxon test confirmed that, overall, H was greater for *GF* (M = 0.845, SD = 0.094) than for LF (M = 0.705, SD = 0.086), z = 24.63, p < 0.001.

Finally, SampEn analysis (Fig. 2, C) revealed a significant effect of force condition, *χ*^2^ *(4)* = 514.21, p < 0.001, Kendall’s W = 0.61, indicating a large effect size. For *GF*, post-hoc Wilcoxon signed-rank tests with Bonferroni correction showed that temporal complexity increased under all constrained conditions (−10%, natural, +10%) compared with the spontaneous pre-condition (all p < 0.001). However, only the -10% condition remained significantly different from the spontaneous post condition (p = 0.003), whereas the natural and +10% conditions did not differ from post (p = 0.104 and p = 0.053, respectively). A significant difference was also found between the spontaneous pre- and post-conditions (p < 0.001), while no differences were observed among the constrained conditions themselves (all p > 0.05). For LF, post-hoc Wilcoxon signed-rank tests showed a more limited pattern. SampEn was significantly higher between the natural and the spontaneous pre-condition (p = 0.016), whereas no other pairwise comparisons reached significance (all p > 0.05). SampEn did not differ among constrained conditions or between spontaneous pre and post trials.

Overall, these results indicate that *GF* is consistently more variable (larger CV), more temporally persistent (larger H) and locally irregular (larger SampEn) than LF. Importantly, *GF* indices were systematically modulated by the constrained nature of the task (−10%, natural, +10% vs. spontaneous pre and post conditions), whereas LF indices remained largely stable across conditions.

### Temporal Evolution of Predictability Within Trials

We analyzed the early and late halves of the hold phases using only H, as this index reflects long range correlation. Figure 3 presents the evolution of H across the first and second halves of each trial, for all five force conditions (spontaneous pre and post, −10%, natural, +10%).

**Figure 3.**
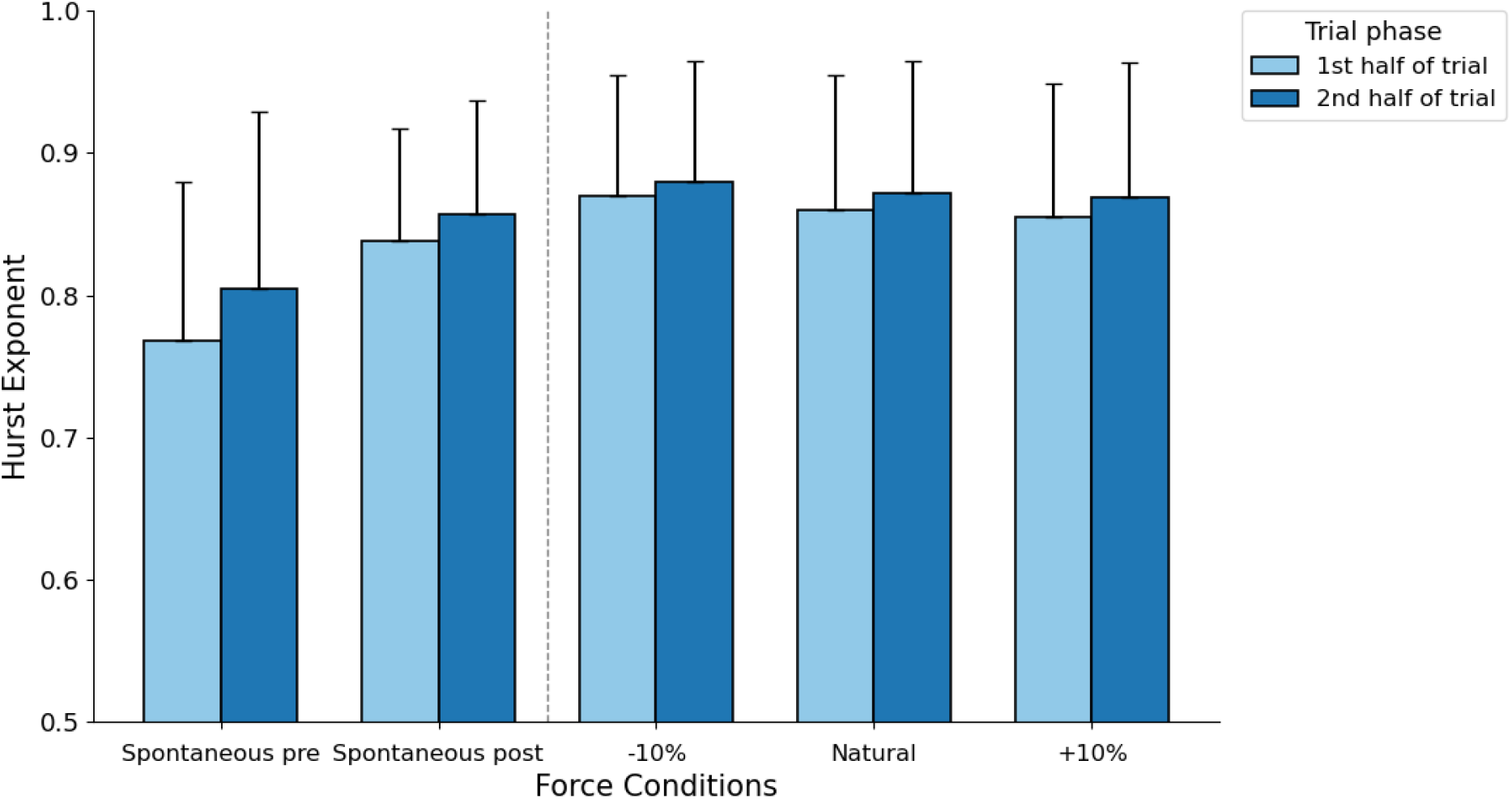
Temporal evolution of GF predictability (H) between the first (light blue) and second halves (dark blue) of the trials across force conditions.

Friedman test revealed a significant main effect of time ( (1) = 87.867, p < 0.001, Kendall’s W = 0.104), indicating that temporal predictability increased throughout the trial. Post-hoc Wilcoxon comparisons confirmed significantly higher H values in the second half (M = 0.857, SD = 0.100) than in the first half (M = 0.838, SD = 0.100; z = -10.633, p < 0.001).

When analyzed per condition, this within-trial increase was pronounced under spontaneous pre and post conditions (both p < 0.001), modest for +10% level (p = 0.04), and non-significant for −10% and natural conditions (p = 0.462, p = 0.235, respectively).

Overall, these results show that the temporal persistence of GF dynamics progressively increases during the trial, especially in self-force contexts, suggesting a greater capacity for adaptive temporal stabilization when the force is not externally constrained.

### Correlations Between Complexity and Variability Indices

We ran a correlation analysis on participant-level mean values that revealed several significant associations between the variability and complexity indices derived from GF and LF signals (Fig. 4).

**Figure 4.**
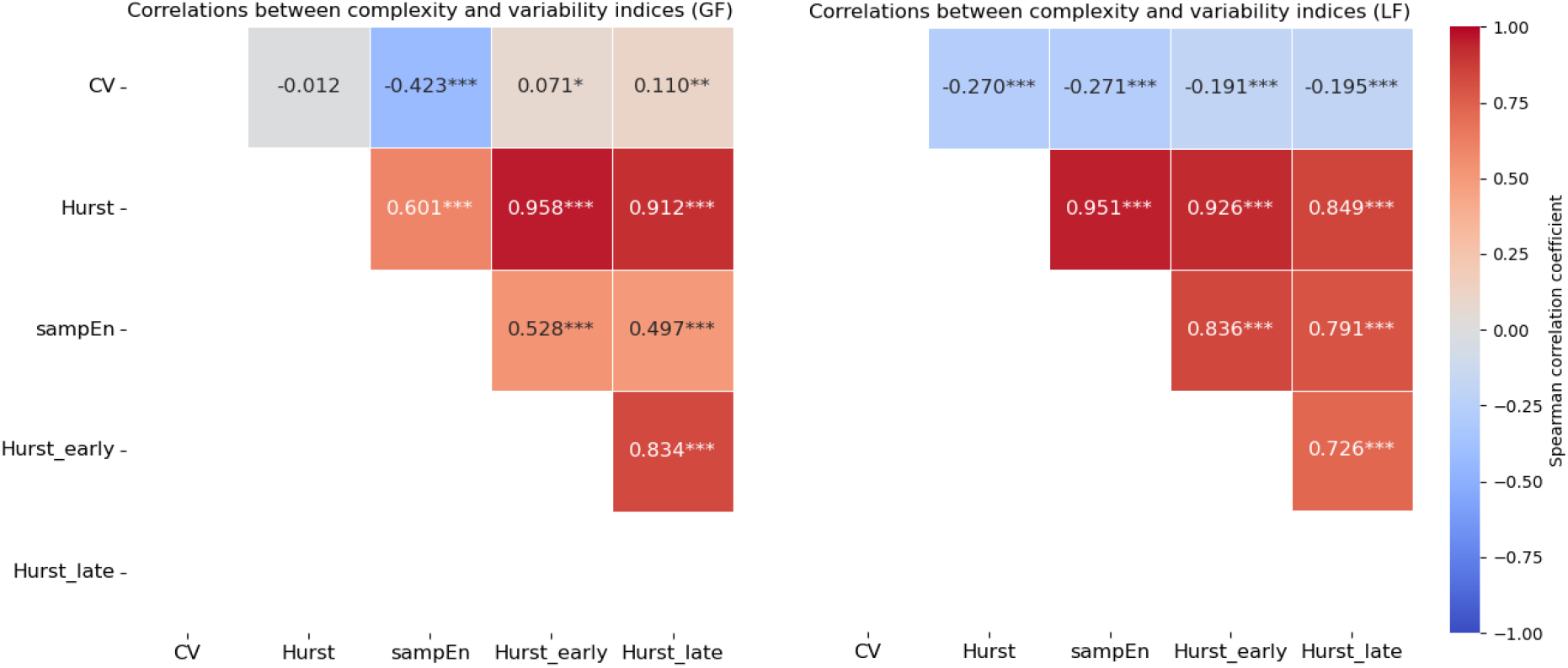
Spearman correlation heatmap between variability (CV) and temporal complexity indices (H and SampEn) extracted from grip force (GF, left panel) and load force (L_F_, right panel). Spearman correlation values are shown and those that were significant are indicated with stars (*p < 0.05, **p < 0.01, ***p <0.001). Strong positive and negative correlations are depicted in dark red and dark blue, respectively. Lighter colors denote weak correlations.

For GF (Fig. 4, left panel), H was strongly and positively correlated with SampEn (ρ = 0.601, p < 0.001), suggesting that signals exhibiting greater temporal predictability tend to be more complex at local time scales. As expected by construction, H also showed very high correlations with its trial-segmented values—both at the beginning (ρ = 0.958, p < 0.001) and end (ρ = 0.912, p < 0.001) of the trial—indicating strong internal consistency in the predictability profile within trials. These two temporal components were themselves significantly correlated (ρ = 0.834, p < 0.001), confirming the intra-trial stability of the H-based measure. Additionally, a moderate negative correlation was found between the CV and SampEn (ρ = −0.423, p < 0.001), indicating that lower amplitude variability is associated with increased local temporal complexity. CV and H were not significantly correlated (ρ = −0.012, p = 0.722), underscoring their complementary roles in characterizing motor variability.

For LF (Fig. 4, right panel), similar trends were observed. H again correlated strongly with SampEn (ρ = 0.926, p < 0.001), as well as with H_early (ρ = 0.951, p < 0.001) and H_late (ρ = 0.849, p < 0.001). The beginning and end segments of the trial were also significantly correlated with each other (ρ = 0.726, p < 0.001), confirming a stable temporal structure in LF as well. In contrast to GF, however, the associations between the CV and the other indices were generally weaker. For example, CV correlated only modestly but significantly with both SampEn (ρ = −0.271, p < 0.001) and H (ρ = −0.270, p < 0.001), suggesting a decoupling between fluctuation magnitude and complexity in LF.

Together, these findings support the view that complexity and temporal predictability are tightly linked within each force level, but that this relationship is stronger in GF than in LF. The CV also quantifies different noise characteristics as compared to H and SampEn. This further reinforces the distinct control strategies deployed by the motor system for these two force components, with GF exhibiting a richer and more structured temporal organization.

## Discussion

The primary objective of this study was to analyze the impact of force constraints of varying nature and intensity on the temporal complexity of human GF. We hypothesized that imposing specific target levels would reduce motor complexity, leading to lower entropy and more rigid temporal dynamics compared with spontaneous grip. Contrary to our initial prediction, we found that target-driven grip force, including the reproduction of the individual natural level, was associated with higher H and higher SampEn values than spontaneous pre-trials. Thus, force constraints did not simply suppress variability; instead, they reorganized it into more temporally persistent yet locally irregular patterns, suggesting an adaptive reconfiguration of motor control under task demands.

### Motor variability and adaptability under constraint

The CVs revealed significantly higher variability amplitude for GF compared to LF. For GF, this variability decreased under all three constrained conditions, whereas it remained stable for LF. This suggests that motor adjustments were primarily expressed through GF. Such a strategy supports models in which behavioral flexibility relies on modulated variability (Faisal et al., 2008; Stergiou et al., 2006). From the perspective of dynamical systems theory, such modulation of variability may represent an adaptive self-organization process where the system explores multiple motor solutions to maintain performance stability (Davids et al., 2003; Seifert et al., 2016). Under constraint, the reduction in variability may reflect a loss of degrees of freedom, consistent with the “Optimal Movement Variability” model proposed by Stergiou and Decker (2011). In this context, such rigidity could be interpreted as a functional adaptation aimed at minimizing errors and optimizing energy cost through tighter motor control.

### Temporal predictability: fractal dynamics altered by constraint

H was consistently higher for GF than for LF, indicating greater temporal predictability in GF. For GF, H increased across all constrained conditions, as well as between the pre and post conditions, confirming its strong sensitivity to dynamic changes. LF remained generally stable, with only a minor effect under the lower constraint condition. These findings are consistent with Fernandes and Chau (2008), who showed that increased motor demands could alter the fractal structure of motor signals.

H reflects the temporal persistence of a signal: H ≈ 0.5 corresponds to random behavior, H > 0.5 indicates persistent trends, and H < 0.5 reflects anti-persistence. In our natural conditions, H ranged between 0.75 and 0.85, indicating persistent but non-saturating long-range temporal dependencies (Goldberger, 1996; Vaillancourt & Newell, 2002). The elevation of H under constraints reflects stronger long-range temporal persistence in the force signal, which may indicate a more constrained mode of control, even if this increased persistence coexists with high local irregularity in the present data.

### Local complexity and entropy: increased information under constraint

SampEn was significantly higher for GF than for LF, indicating a greater informational richness in the GF signal. It increased significantly under certain constraints, particularly between the pre and post conditions. For LF, no significant variation was observed. The LF signal results from the addition of a gravitational term (mg) and an inertial term (ma) (White, 2015). The first is largely determined by physical constraints, including object mass and gravitational acceleration. The second depends on the acceleration which had to be minimal in this task, since the object had to be stabilized. LF may therefore carry less task-related variability than GF in this static context. These results suggest that some forms of constraint enhance the adaptability of the neuromotor system, reflected by increased local complexity, particularly in GF.

High SampEn values reflect greater signal irregularity and reduced predictability. While theoretical simulations on fractal processes typically demonstrate an inverse relationship where increased self-similarity (H) leads to reduced entropy (Omidvarnia et al., 2018), our results reveal a paradoxical dynamic. In the present data, this increase coincided with a simultaneous rise in H.

Contrary to classical optimal variability frameworks–which suggest that increased predictability implies reduced local complexity–our findings indicate that global temporal organization and local informational richness can coexist. This suggests that motor control may combine temporal stability with local flexibility, inviting a more nuanced view of control than a simple dichotomy between rigid and complex control.

### Inter-trial dynamics: adaptive stability or reset effect?

Inter-trial analyses revealed stable variability and complexity indices from one trial to the next, with no significant differences within each condition. This lack of change may be explained by three factors: (1) the randomization of trials, which imposed a different constraint each time, likely forcing a constant motor “reset”; (2) sustained attentional engagement due to visual feedback, which prevented automatization; and (3) greater motor freedom in the natural condition, allowing the system to adjust without altering its overall complexity.

The remarkable stability of LF, both in mean values and variability, confirms that it is the variable being stabilized by the system. This also indicates that GF serves as the regulatory variable, with greater flexibility and room for adjustment. This differentiated organization of degrees of freedom supports models suggesting that motor control is strategically adapted to the task’s specific goals. It should also be noted that constrained and spontaneous conditions differed in the presence of a visual force target. This coupling is task-intrinsic: the target cursor was the operational instantiation of the constraint, not an independent variable layered onto a common task. Thus, the relevant distinction is between two motor tasks: spontaneous force maintenance and online error correction against an explicit reference.

### Intra-trial dynamics: progressive adjustments and evolving complexity

The intra-trial analysis, which examines the evolution of fractal indices within a single trial, revealed progressive temporal adjustments for GF. In natural conditions, H increased significantly from the beginning to the end of the trial, reflecting a process of self-organization—the system refines its strategy over the course of the task. This illustrates adaptive motor control, consistent with previous findings on motor stabilization.

In contrast, under constrained conditions, this evolution was either absent or attenuated, except in the +10% condition (p = 0.04). In this case, the progressive adjustment may be due to an increased demand for precision beyond the usual safety margin. The system, already highly engaged from the start, appears to adopt a rigid control strategy. The delayed increase in H may reflect a compensatory adaptation to increased task demands. These findings suggest that the freedom to explore, present in natural conditions, is essential for the motor system to dynamically adjust its strategy over time.

Thus, intra-trial dynamics reveal not only the system’s initial state but also its capacity for real-time reorganization. They enrich the interpretation of motor complexity by distinguishing an adaptive system from one that becomes fixed under constraint.

### Correlations between variables: an integrated view

The strong positive correlations between H and SampEn, particularly under constrained conditions, suggest that high temporal predictability coexists with high local complexity. This positive association suggests that, at least for continuous force control in healthy adults, increased temporal predictability does not necessarily come at the expense of local informational richness. Instead, the neuromotor system can simultaneously exploit long-range correlations and short-range irregularity, pointing to a more nuanced notion of “optimal complexity” than a simple inverse mapping between order and variability.

Far from contradicting classical models of optimal variability, such as that of Stergiou (Stergiou et al., 2006)—which rely primarily on spatial measures of complexity (e.g., fractal dimension)—these results may be viewed as a complementary extension, highlighting the richness of temporal dynamics in the study of GF. The co-increase of H and SampEn observed here suggests that a structured motor system can preserve local informational richness, reflecting a healthy dynamic that combines global stability with local flexibility. These findings support a broader conceptualization of motor complexity, in which long-range temporal persistence, local irregularity, and variability amplitude are viewed as complementary rather than redundant dimensions of motor control.

In contrast, correlations between H or SampEn and the CV were weak or absent. This suggests that the amplitude of variability (CV) is independent from its temporal structure. A signal may thus exhibit large variability without being complex (e.g., pathological tremor), or, conversely, be minimally variable yet highly complex (e.g., the invisible postural adjustments of a tightrope walker). This dissociation reinforces the value of a multi-indicator approach that combines amplitude (CV), temporal persistence (H), and local complexity (SampEn) to better capture the richness of motor control. Such an integrated perspective is particularly relevant for detecting subtle neuromotor adjustments in response to task constraints and may inform future clinical assessments or targeted training approaches (Manor & Lipsitz, 2013). Under constraint, the system appears to restrict the amplitude of fluctuations while enhancing both long-range temporal structure and local irregularity. This pattern is compatible with a strategy where variability is not eliminated but redistributed across scales to maintain adaptability while satisfying accuracy demands. Thus, analyzing the structure of variability—beyond its mere amplitude—provides deeper insight into the adaptive mechanisms of the motor system, especially under constraint, where fractal reorganization strategies appear to take precedence over simple amplitude-based adjustments.

### Limitations of the Study

Despite the relevance of the findings, several limitations should be acknowledged. The small sample size (n = 20) limits the generalizability of the conclusions, particularly regarding interindividual variability. A larger and more diverse population (in terms of age, sex, and expertise) would help refine the interpretation. Furthermore, fractal analysis using H and SampEn, while informative, remains sensitive to several parameters (e.g., windowing, thresholds, measurement noise), which may affect the reliability of the indices. The inclusion of additional nonlinear indices, such as multiscale entropy or Lyapunov-based measures, could further enrich the approach. Moreover, the pre/post design, even when interspersed with rest periods, may introduce a bias in the absence of direct fatigue measurements. Despite randomization of the constrained trials, potential learning or fatigue effects cannot be entirely ruled out.

Future studies could further probe the respective contributions of target-driven regulation and visual display by independently manipulating these factors in designs where both can be independently varied. Furthermore, this study focused on the dominant hand in a static task. Comparisons with the non-dominant hand, older adults, clinical populations (e.g., stroke, Parkinson’s disease), or dynamic tasks would help test the robustness of these markers in more ecological and varied contexts.

Finally, both H and SampEn assume quasi-stationary signals over the analyzed segments. Although we minimized non-stationarities by removing grasp and release transients and selecting stable hold phases, residual trends or slow drifts may still affect the estimates and should be considered when generalizing the present findings.

## Conclusion

In summary, our findings demonstrate that imposing force constraints modifies the structure of motor variability, particularly in GF. The results show that temporal complexity indices, both fractal (H) and entropic (SampEn), vary significantly depending on the imposed motor constraints, reflecting dynamic adjustments in motor control. The observed reduction in variability amplitude under constraint does not reflect a simpler or less adaptable form of control, but rather a restructuring of variability across scales, combining increased temporal persistence with greater local irregularity.

However, the positive covariation between H and SampEn observed here adds nuance to current theoretical models by showing that a motor system can simultaneously exhibit temporal stability and local complexity. These observations reinforce the value of a multidimensional assessment of motor complexity that jointly considers persistence, local irregularity, and variability amplitude. Ultimately, this approach may help refine clinical assessment and rehabilitation strategies by identifying variability profiles characteristic of either healthy or pathological motor function. Together, these findings advance theoretical understanding of how the motor system organizes variability under constraint: rather than suppressing complexity, the system redistributes it across temporal scales, combining increased long-range persistence with preserved local irregularity. This multidimensional view of motor complexity refines current theoretical frameworks and provides new markers for assessing adaptive capacity in both healthy and clinical populations.

